# GLUT4 expression and glucose transport in human induced pluripotent stem cell-derived cardiomyocytes

**DOI:** 10.1101/646828

**Authors:** Peter R.T Bowman, Godfrey L. Smith, Gwyn W. Gould

**Affiliations:** Henry Wellcome Laboratory of Cell Biology, Institute of Molecular Cell and Systems Biology, Davidson Building, College of Medical, Veterinary and Life Sciences, University of Glasgow, Glasgow G12 8QQ. United Kingdom; Institute of Cardiovascular and Medical Sciences, College of Medical, Veterinary and Life Sciences, University of Glasgow, Glasgow G12 8QQ. United Kingdom

## Abstract

Induced pluripotent stem cell derived cardiomyocytes (iPSC-CM) have the potential to transform regenerative cardiac medicine and the modelling of cardiac disease. This is of particular importance in the context of diabetic cardiomyopathy where diabetic individuals exhibit reduced cardiac diastolic contractile performance in the absence of vascular disease, significantly contributing towards high cardiovascular morbidity. In this study, the capacity of iPSC-CM to act as a novel cellular model of cardiomyocytes was assessed. The diabetic phenotype is characterised by insulin resistance, therefore there was a specific focus upon metabolic parameters. Despite expressing crucial insulin signalling intermediates and relevant trafficking proteins, it was identified that iPSC-CM do not exhibit insulin-stimulated glucose uptake. iPSC-CM are spontaneously contractile however contraction mediated uptake was not found to mask any insulin response. The fundamental limitation identified in these cells was a critical lack of expression of the insulin sensitive glucose transporter GLUT4. Using comparative immunoblot analysis and the GLUT-selective inhibitor BAY-876 to quantify expression of these transporters, we show that iPSC-CM express high levels of GLUT1 and low levels of GLUT4 compared to primary cardiomyocytes and cultured adipocytes. Interventions to overcome this limitation were unsuccessful. We suggest that the utility of iPSC-CMs to study cardiac metabolic disorders may be limited by their apparent foetal-like phenotype.

## Introduction

Diabetes is one of the leading healthcare challenges worldwide. Whilst the most common major complication of this condition is vascular disease and therefore increased incidence of stroke or myocardial infarction, there is also a significantly elevated direct risk of heart failure [1]. This is due in part to an impairment of diastolic cardiac contractile function in diabetic individuals independent of vascular disease termed diabetic cardiomyopathy (DCM) [2–4]. Given the high rate of cardiovascular mortality associated with diabetes combined with the lack of DCM specific treatments available, improved understanding of the pathophysiological mechanisms underlying this condition is of clinical relevance.

As DCM progresses over time, structural changes such as concentric hypertrophy and fibrosis are observable [5,6] and will undoubtedly contribute to further impairments in cardiac function. However, there is evidence from human populations that reduced cardiac contractility is observable prior to the onset of structural remodelling [7], suggesting that this is a (maladaptive) response to - rather than central causal factor of - this condition. Accordingly, the original physiological deficit underlying DCM likely originates from within the individual contractile units of the heart – the cardiomyocytes.

It is challenging to obtain information regarding this pathophysiological mechanism from human samples. However, the *db/db* mouse model captures several clinical features of DCM such as weight gain and impaired cardiac function in the presence of hyperglycemia and hyperinsulinemia but absence of atherosclerosis or hypertension [8]. Impaired cardiac contractile function has been recorded from these mice through *in vivo* echocardiography (impaired ejection fraction and E/A ratio), *ex vivo* Langendorff working heart preparations (reduced rate of LV pressure development and decay), and individual isolated cardiomyocytes (impaired fractional shortening and rate of relaxation) [9,10]. These studies identified deficits in intracellular calcium handling as the fundamental mechanism underlying this impaired contractile activity.

In healthy muscle and adipose tissue, activation of the insulin receptor in turn activates signalling cascades that ultimately initiate the translocation of the glucose transporter GLUT4 from a specialised intracellular retention site to the plasma membrane [11]. A defining feature of the (type II) diabetic phenotype is the presence of peripheral insulin resistance (IR), whereby this action of insulin is impaired. There is evidence to suggest that reduced glucose uptake coupled with coinciding increased fatty acid uptake and oxidation in the diabetic heart precedes and possibly contributes towards impaired cardiomyocyte calcium handling/contractility. The oxidation of fat is associated with an increased oxygen cost and in line with this reduced cardiac efficiency has been recorded from diabetic individuals [12], thereby placing increased strain upon the heart. Additionally, the intracellular PCr/ATP ratio was observed to be reduced in human diabetic myocardium [13]. This would be expected to impair the high energy phosphate shuttle system and is linked to a reduction in maximum workload capacity [14]. Combined, these mechanisms may create a large mismatch between ATP availability and demand, and therefore limit the ATPase action of proteins including SERCA and myosin. In combination this could account for impaired contractility in DCM. In support of this, overexpression of GLUT4 in the *db/db* mouse model normalises the observed aberrant metabolic and contractile cardiac phenotype [15,16].

Many studies suggest that insulin signalling and/or glucose metabolism (or a lack thereof) have important roles in the development of DCM. For example, insulin signalling may be required to reduce mitochondrial uncoupling and enable economical oxidation of fatty acids [17], and the more oxygen efficient oxidation of glucose may be essential under periods of subclinical myocardial stress, which likely precede DCM. Although the use of animal-based DCM models has facilitated significant progress in this area of research, human studies are necessary to fully understand these mechanisms to the extent that effective interventions can be developed. The generation of human induced pluripotent stem cell derived cardiomyocytes (iPSC-CM) was first published in 2007 [18] and could be of relevance to this field. iPSC-CM have gained attention due to their potential to act as both the basis of regenerative medicine interventions in the context of myocardial infarction, and also as a novel cellular model for different disease phenotypes, including DCM. Whilst these cells exhibit cardiomyocyte specific gene/protein expression and general function, it is widely accepted that they correspond to foetal-like cardiomyocytes with regards to their electrophysiology, structure, and calcium handling [19,20]. Particularly apparent limitations are their small circular phenotype and lack of t-tubules, which results in non-uniform intracellular calcium release across the width of the cells and contributes to their relatively weak contractile output [21,22].

It has been shown that through culturing iPSC-CM in medium designed to replicate the blood chemistry of a diabetic individual a phenotype corresponding to DCM was induced in iPSC-CMs, based upon the onset of cellular hypertrophy, reduced frequency of calcium transients, and an increased irregularity of contractile rate [23]. However, these studies did not examine the glucose transport or insulin-sensitivity of iPSC-CMs, and therefore could not determine if these cells exhibited (or were even capable of exhibiting) insulin resistance. Whilst prior work has demonstrated that under standard high glucose cell culture conditions these cells rely primarily upon glycolysis to generate ATP, they have been shown to have a degree of metabolic flexibility [24]. However, there is no evidence to indicate whether or not iPSC-CM exhibit insulin stimulated glucose transport or GLUT4 trafficking, a defining feature of mature insulin sensitive cardiomyocytes. Therefore, the key aim of this study was to investigate whether iPSC-CM exhibit insulin-stimulated glucose uptake in order to assess their potential to act as the basis of a novel cellular model of DCM.

## Materials and Methods

### Cell Culture

iPSC-CM were commercially obtained directly from Axiogenesis [note this company has recently merged with Pluriomics to form NCardia] (#Ax-B-HC02-1M) and Cellular Dynamics International (CDI; #CMC-100-012-000.5). Upon receipt, cells were transferred to liquid nitrogen for long term storage. Thereafter, when required for use, cells were retrieved and plated according to manufacturer’s instructions into wells of a 96-well plate coated with fibronectin from bovine plasma (Sigma) at a final concentration of ~2 μg/cm^2^. Axiogenesis and CDI iPSC-CM were plated as recommended at a density of 35,000 or 50,000 viable cells per well respectively, unless stated otherwise. Cells were maintained in a sterile humidified incubator (37°C, 5% CO_2_) in the appropriate maintenance medium provided by the manufacturers, and was replaced 4-24 hours post plating, and every 24-48 hours thereafter.

3T3-L1 fibroblasts were obtained directly from ATCC (#CL-173) and grown and differentiated as outlined previously [25].

### Primary rabbit cardiomyocyte isolation

Procedures were undertaken in accordance with the United Kingdom Animals (Scientific Procedures) Act of 1986 under a project licence (PPL70/8835) and conform to the Guide for the Care and Use of Laboratory Animals published by the US National Institutes of Health and approved by the Glasgow University Ethical Review Board. White rabbits (20 weeks old; 3.0–3.5 kg) were euthanized with an intravenous injection of sodium pentobarbital (100 mg/kg) and 2500 IU heparin. Excised hearts were reverse perfused on a Langendorff rig with a Krebs buffer supplemented with 5000 U/mL heparin. Subsequently, protease (33 μg/mL) and collagenase (217 IU/mL) were added to this solution for approximately 15 min, prior to manual mincing with a sterile surgical scalpel blade. Finally, the solution was filtered to remove unprocessed tissue. Where required, cell lysates were generated by pelleting cells and then resuspending them in RIPA buffer. Cells were maintained on ice for 20 min in between 2 rounds of manual homogenisation via a Dounce style tissue grinder. Thereafter lysates were centrifuged at 17,500 x g for 15 min at 4°C and the supernatant was collected.

When used for measurement of glucose uptake, cardiomyocytes were resuspended in several cycles of Krebs buffer (130 mM NaCl, 5 mM Hepes, 4.5 mM KCl, 0.4 mM NaH_2_PO_4_, 3.5 mM MgCl_2_, 10 mM glucose, 140 μM CaCl_2_) supplemented with increasing concentrations of calcium chloride, until a final concentration of 1.8 mM was attained. Thereafter cells were plated at 15,000 viable cells per well of 96-well plates, maintained in Medium 199 (1 mM L-glutamine, 5 mM creatine, 5 mM taurine, 5 mM carnitine), and assayed approximately 3 hours post-plating.

### Generation of primary mouse cardiomyocyte lysates

Frozen segments of myocardial tissue from 20 week old male 129/Sv mice were kindly donated by Dr Anna White. These were then manually diced in RIPA buffer prior to 2 rounds of homogenisation (with 20 min break) using a Dounce style tissue grinder, whilst being kept on ice at all times. Lysates were then centrifuged at 17,500 x g for 15 min at 4°C and the supernatant was collected. Due to the low output obtained, lysates from 2-4 individual hearts were pooled to form each of the samples used for subsequent immunoblotting.

### SDS-PAGE and Immunoblotting

iPSC-CM and 3T3-L1 adipocyte lysates were generated directly via application of 2x Laemmli sample buffer (LSB) for 20 min whilst plates were maintained on ice. Thereafter, the samples were collected and heated to 65°C for 10 min prior to immediate use, or temporary storage at −20°C. Prior to immunoblotting with primary mouse cardiomyocyte lysate, samples were thawed on ice, combined 1:1 with 2x LSB, and heated to 65°C for 10 min. Samples were separated on acrylamide gels and immunoblotting performed as described previously [25]. Quantification of target protein expression was performed via densitometry and normalised against GAPDH.

### Antibodies

Anti-pan Akt (#2920), anti-phospho S473 Akt (#4058), anti-ERK1/2 (#9102), anti-phospho ERK1/2 (#9106) were from Cell Signalling (Danvers, Massachusetts, USA). Anti-Sx4 (#110042) and Anti-SNAP23 (#111202) were from Synaptic Systems (Goettingen, Germany). Anti-GLUT1 (#652) and anti-GLUT4 (654) were from AbCam (Cambridge, United Kingdom) and anti-GAPDH (#4300) was from Ambion (Foster city, California, USA). Detection antibodies were from LI-COR Biosciences (Lincoln, Nebraska, USA).

### [^3^H]-2-deoxyglucose uptake assay

Prior to performing this assay 3T3-L1 adipocytes and iPSC-CM on 96-well plates were incubated in serum free medium (DMEM) for 2-4 hours. Cells were then transferred onto hotplates (maintained at 37°C) and washed twice in Krebs Ringer Phosphate (KRP) buffer (128 mM NaCl, 4.7 mM KCl, 5 mM NaH_2_PO_4_, 1.25 mM MgSO_4_, 1.25 mM CaCl_2_). Subsequently, cells were maintained in KRP +/− 860 nM insulin for 20 min, prior to the addition of [^3^H]-2-deoxyglucose (50 μM 2-deoxyglucose and 0.4 μCi [^3^H]-2-deoxyglucose). Parallel incubations in the presence of 40 μM cytochalasin B were performed to account for non-specific association of isotope with the cells. All values reported have been corrected for this.

For assays using the selective GLUT inhibitor BAY-876 [26] (Sigma), cells were incubated with this compound at the concentrations shown for 20 min prior to the addition of the assay mix. Where iPSC-CM were treated with blebbistatin (AbCam), which prevents myocyte contractile activity via inhibition of Myosin ATPase activity, cells were treated with the specified concentration for approximately 3 hours prior to and throughout the assay.

### Statistical analysis

Statistical testing was performed with GraphPad Prism 7. Where appropriate, the relevant statistical test that was implemented is reported, but in general this was an unpaired t-test, or a 1 or 2 way ANOVA. The level of significance was set at P=0.05.

## Results

### iPSC-CM express core elements required for insulin stimulated glucose uptake

In order to gain an overview of the machinery present in iPSC-CM, immunoblotting was used to determine the presence or absence of proteins that are essential in mediating insulin stimulated GLUT4 trafficking. As can be observed in Fig 1, the insulin signalling intermediates Akt (Protein kinase B) and ERK1/2 (MAPK 42/44) were expressed and both capable of exhibiting insulin-stimulated phosphorylation. Akt is a central factor in insulin-stimulated glucose transport [11], and there is strong evidence implicating ERK1/2 kinases in the regulation of glucose uptake in human muscle [27]. Additionally, both of the plasma membrane t-SNAREs known to mediate the fusion of GLUT4-containing vesicles with the surface of adipocytes and myocytes, Syntaxin 4 and SNAP23, are expressed in these cells (data not shown).

**Fig 1.**
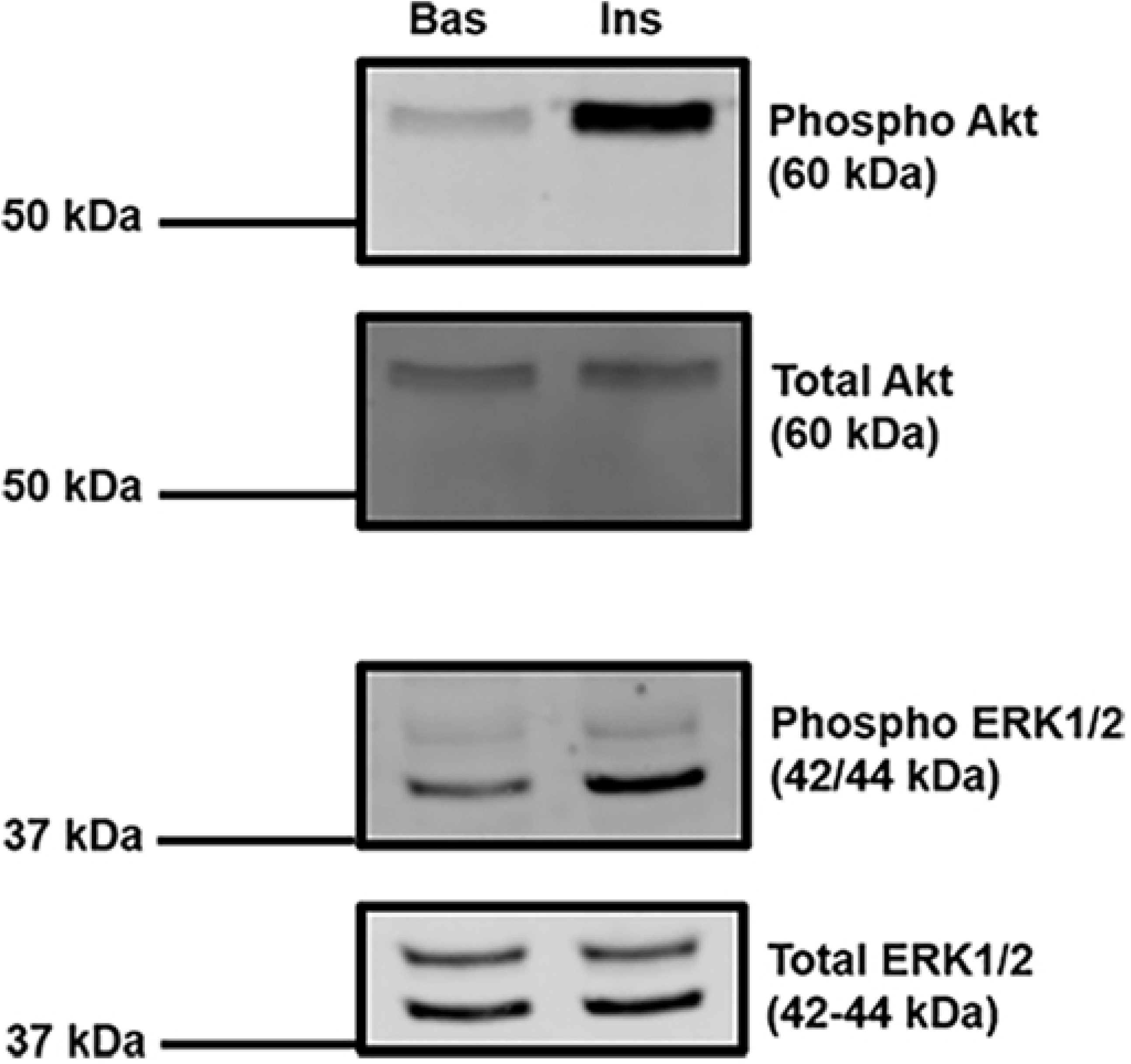
Key insulin signalling intermediates are present in iPSC-CM. Protein lysates were generated from iPSC-CM and subjected to SDS-PAGE and immunoblotting. Lysates were incubated with antibodies probing for the expression of total ERK1/2 (1:2000, 3% BSA, TBS-T), phospho-ERK1/2 (1:1000, 3% BSA, TBS-T), total Akt (1:2000, 3% BSA, TBS-T), and phospho-Akt (ser473, 1:1000, 3% BSA, TBS-T). For Akt, lysate generated from approximately 50,000 CDI iPSC-CM was loaded per lane. This experiment was repeated in Axiogenesis iPSC-CM, with at least 3 independent replicates for each cell type. For ERK1/2, lysate generated from approximately 20,000 Axiogenesis iPSC-CM was loaded per lane. This experiment was performed with Axiogenesis iPSC-CM only, and repeated with 3 independent samples. Where indicated cells were stimulated with 860 nM porcine insulin for 30 min prior to lysis. The approximate position of molecular weight markers are indicated. Bas = basal, unstimulated; Ins = insulin stimulated.

### iPSC-CM do not exhibit insulin stimulated glucose uptake

2-deoxy-D-glucose uptake assays were optimised for cells on a 96-well plate format. As shown in Fig. 2, a clear basal signal and subsequent insulin response could be detected from both 3T3-L1 adipocytes and isolated primary rabbit cardiomyocytes. However, insulin failed to significantly increase glucose uptake in iPSC-CM (Fig. 2). The data presented is from a single commercial source of cells, however similar results were obtained from iPSC-CM from a second independent supplier (Supplemental fig 1).

**Fig 2.**
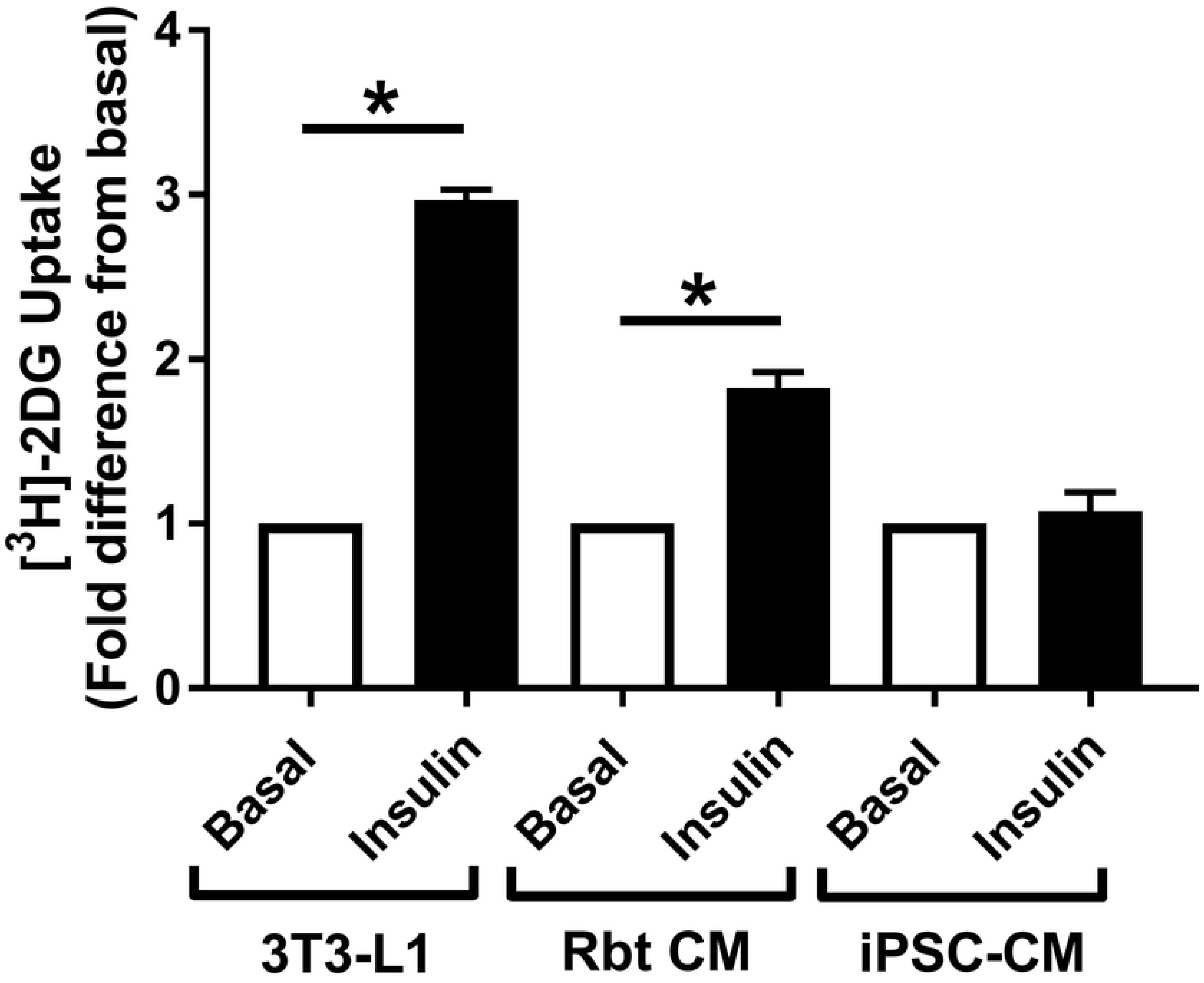
Insulin stimulated [^3^H]-2-deoxyglucose uptake in iPSC-CM, 3T3-L1 adipocytes, and primary adult cardiomyocytes. Background corrected [^3^H]-2-deoxyglucose uptake is displayed from several cell types, generated via the protocol detailed in the methods. 3T3-L1 adipocytes and CDI iPSC-CM were serum starved for 4 hours prior to the assay, insulin stimulated for 20 min, and incubated with [^3^H]-2-deoxyglucose assay mix for 10 min. Primary adult rabbit septal cardiomyocytes were assayed 3 hours post plating, insulin stimulated for 20 min, and incubated with [^3^H]-2-deoxyglucose assay mix for 20 min. Data is displayed as the mean (+ S.E.M) fold change in uptake relative to basal values from 3 (iPSC-CM and 3T3-L1 adipocytes) or 7 (cardiomyocytes) individual experiments. Basal values corresponded to an average of 1311 CPM (rabbit cardiomyocytes), 3167 CPM (3T3-L1 adipocytes), or 2229 CPM (iPSC-CM) respectively, per well of a 96-well plate. Statistical testing was performed with an unpaired t-test on raw unadjusted data. * indicates P<0.05.

### Contraction is a key regulator of glucose uptake in iPSC-CM

In addition to insulin, contraction is a potent stimulus to initiate GLUT4 translocation to the plasma membrane and therefore also increase cellular glucose uptake [28]. Whilst insulin-stimulated glucose uptake can be recorded in contracting myocardium, this response is considerably less powerful than that recorded from quiescent isolated cardiomyocytes [29,30]. It was considered possible that the spontaneous contractile activity of iPSC-CM could potentially be masking a small yet significant insulin response. In order to assess this possibility basal and insulin stimulated glucose uptake were recorded in iPSC-CM in the presence and absence of blebbistatin. Inhibition of contraction via this method resulted in a significant reduction in iPSC-CM glucose uptake by >50%, in the presence or absence of insulin (Fig. 3). However, regardless of whether cells were contracting or not, insulin failed to significantly increase glucose uptake. This suggests that whilst contraction is undoubtedly a critical regulator of metabolic demand and therefore substrate uptake in iPSC-CM, it does not interfere with any metabolic regulation of these cells by insulin.

**Fig 3.**
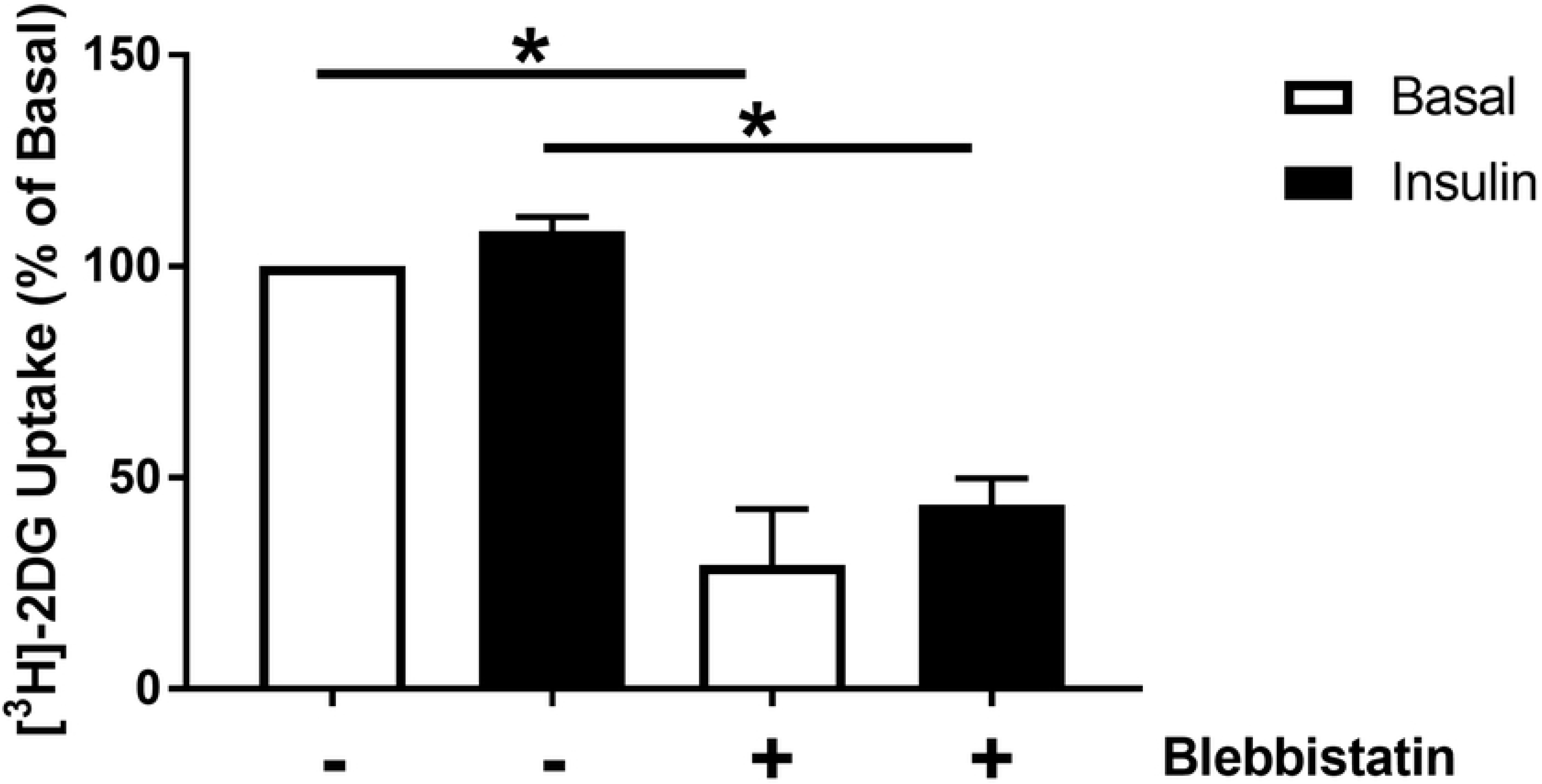
Glucose uptake in quiescent iPSC-CM. Cells were incubated with 3-10 μM blebbistatin for 3 hours prior to and throughout the measurement of [^3^H]-2-deoxyglucose uptake in iPSC-CM. Cells were stimulated with 860 nM porcine insulin prior to incubation with [^3^H]-2-deoxyglucose assay mix. Data is displayed as the mean (+S.E.M.) percentage difference in uptake relative to basal values from 3 individual experiments. Statistical testing was performed with a 2-way ANOVA on raw unadjusted data. * indicates P<0.05.

### iPSC-CM do not express GLUT4

Whilst there are numerous proteins and interactions responsible for the regulated trafficking of GLUT4, ultimately one of the foundational requirements to facilitate this process is sufficient expression of GLUT4 itself. *In vivo*, cardiac GLUT1 expression predominates during early development prior to a metabolic switch and rapid induction of GLUT4 protein during early post-natal life with a corresponding repression of GLUT1 [31]. Given the generally reported classification of iPSC-CM as foetal like cardiomyocytes due to aspects of their electrophysiological and structural phenotype, it could be that these cells express the non-insulin sensitive GLUT1 transporter in favour of GLUT4.

Therefore, quantification of GLUT1 and GLUT4 was performed in iPSC-CM and the primary adult cardiomyocytes they are claimed to represent, by using 3T3-L1 adipocytes as a reference. Quantified expression of each transporter in this cell type has been published previously, with an estimated 950,000 and 280,000 copies of Glut 1 and 4, respectively, per cell [32]. As shown in Fig. 4 and Table 1, by comparing the expression of GLUT4 normalised against GAPDH (loading control) in each source of cardiomyocytes against the signals obtained from 3T3-L1 adipocytes, our data suggest that iPSC-CM express approximately 8-fold less GLUT4 than primary adult cardiomyocytes. In contrast, completion of identical analysis for GLUT1 (Fig. 5) revealed iPSC-CM to express approximately 30-fold more of this transporter, relative to primary cardiomyocytes.

**Fig 4.**
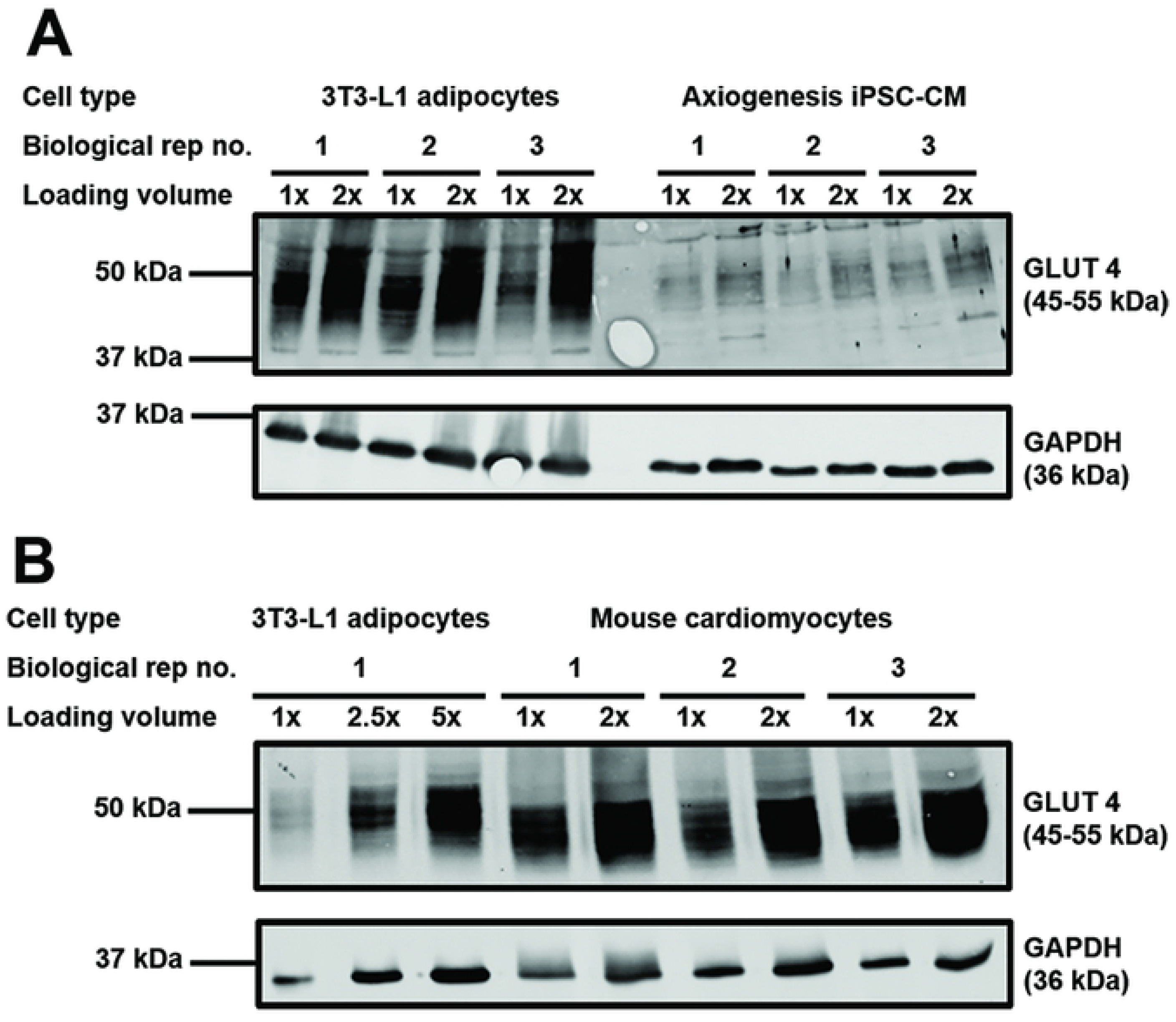
GLUT4 protein expression in different cardiomyocyte models. Protein lysates generated from 3T3-L1 adipocytes, primary adult mouse cardiomyocytes, and iPSC-CM were subjected to SDS-PAGE and immunoblotting. Lysates were incubated with antibodies probing for GLUT4 (1:2000, 1% milk, PBS-T) and GAPDH (1:80,000, 1% milk, PBS-T). Two different amounts of each sample of cardiomyocyte lysate from three biologically independent sources were loaded, in addition to at least three samples of 3T3-L1 adipocyte lysate. For each cell type, 1x refers to lysate generated from approximately 15,000 (3T3-L1 adipocytes) or 17,500 (iPSC-CM) cells, or approximately 20 μg of protein (mouse cardiomyocytes). Panel A compares 3T3-L1 adipocytes and iPSC-CMs; Panel B compares 3T3-L1 adipocytes and primary mouse cardiomyocytes.

**Fig 5.**
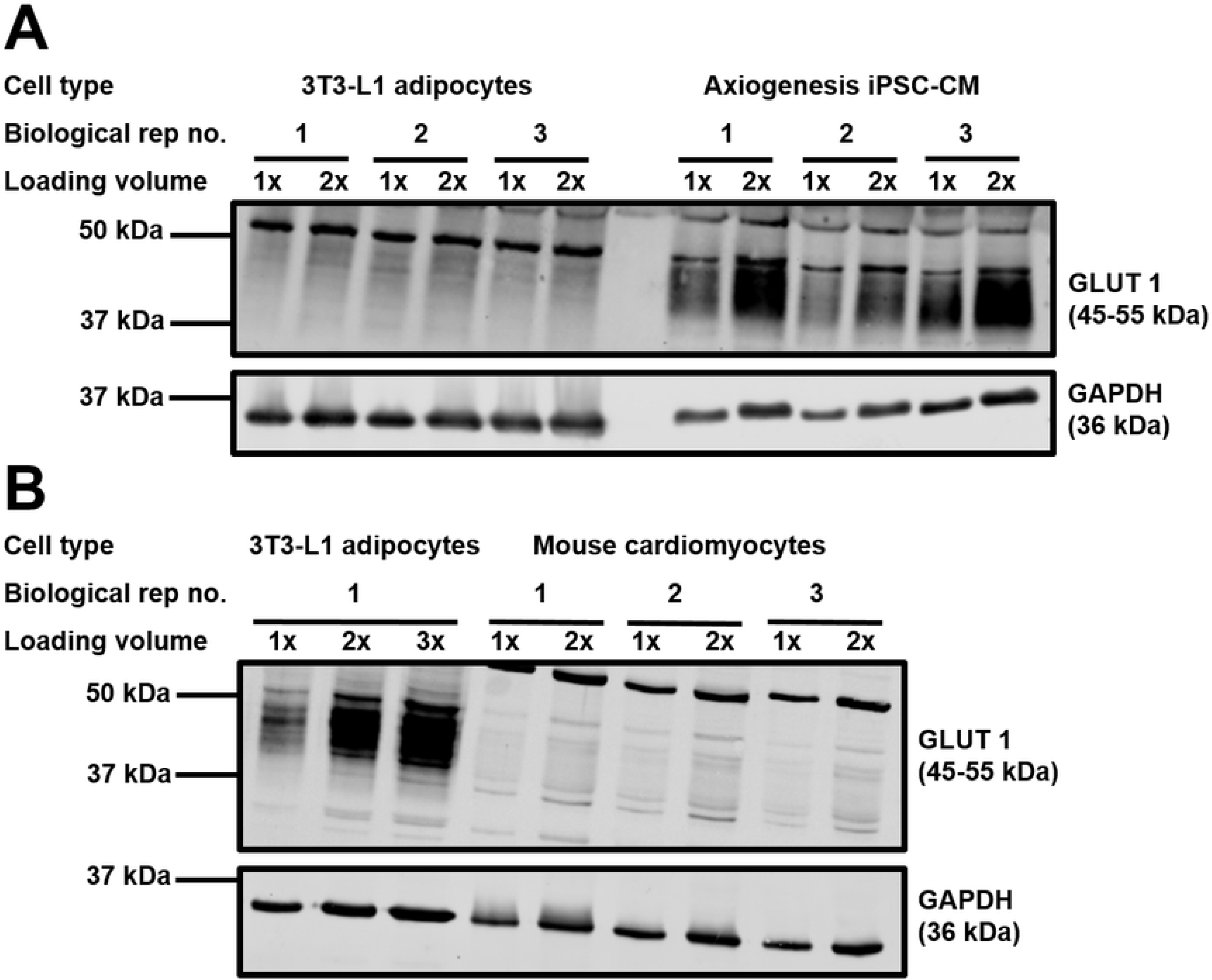
GLUT1 protein expression in different cardiomyocyte models. Protein lysates generated from 3T3-L1 adipocytes, primary adult mouse cardiomyocytes, and iPSC-CM were subjected to SDS-PAGE and immunoblotting. Lysates were incubated with antibodies probing for GLUT1 (1:1000, 1% milk, PBS-T) and GAPDH (1:80,000, 1% milk, PBS-T). Two different amounts of each sample of cardiomyocyte lysate from three biologically independent sources were loaded, in addition to at least three samples of 3T3-L1 adipocyte lysate. For each cell type, 1x refers to lysate generated from approximately 15,000 (3T3-L1 adipocytes panel A), 17,500 (iPSC-CM), or 45,000 (3T3-L1 adipocytes panel B) cells, or approximately 20 μg of protein (mouse cardiomyocytes). Panel A compares 3T3-L1 adipocytes and iPSC-CMs; Panel B compares 3T3-L1 adipocytes and primary mouse cardiomyocytes.

**Table 1.** Quantification of GLUT1 and GLUT4 protein expression in different cardiomyocyte models.

| Cell Type | Ave. Fold Difference GLUT1 | Estimated Amount GLUT1 (ng/1.5 mg total protein) | Ave. Fold Difference GLUT4 | Estimated Amount GLUT4 (ng/1.5 mg total protein) |
| --- | --- | --- | --- | --- |
| 3T3-L1 Adipocytes | 1 | 215 | 1 | 45 |
| iPSC-CM | 5.1 fold higher | 1106 | 3.1 fold lower | 14.6 |
| Primary<br>Mouse CM | 5.7 fold lower | 37.5 | 2.7 fold higher | 121.4 |

Quantification of target protein expression was performed via densitometry and normalised against GAPDH to account for differences in loading. Estimates for differences in absolute GLUT expression were generated by calculation of the fold difference between cardiomyocyte lysates and 3T3-L1 adipocytes across 2 replicate immunoblots.

In order to gain further insight beyond this protein expression data, the effect of GLUT1 and GLUT4 inhibition upon iPSC-CM glucose transport was assessed. In order to achieve this the drug BAY-876 was utilised. BAY-876 inhibits GLUT1 with an IC50 of 2 nM, and inhibits GLUT4 with an IC50 of 200 nM [26]. In three independent experiments, BAY-876 inhibition of GLUT1 (20 nM) significantly (P<0.05) reduced [^3^H]-2-deoxyglucose uptake into 3T3-L1 adipocytes in the presence or absence of insulin by approximately 25% (Fig. 6). Increasing the concentration of BAY-876 to 2 μM resulted in a further significant (P<0.05) reduction in uptake values to almost no detectable signal. This strongly suggests that in this cell type GLUT4 is predominantly responsible for regulating glucose uptake, both under basal and insulin stimulated conditions. It is particularly striking that a significant insulin response was only not recorded (P>0.05) in the presence of 2 μM BAY-876. Inhibition of the action of GLUT1 in iPSC-CM produced a larger (~50%) significant (P<0.05) decrease in [^3^H]-2-deoxyglucose uptake (Fig. 6). Furthermore, in contrast to 3T3-L1 adipocytes, increasing the concentration of BAY-876 to 2 μM had no significant (P>0.05) additional effect upon transport values. These results indicate that iPSC-CM do not rely upon GLUT4 for the regulation of cellular glucose uptake and are more heavily dependent upon GLUT1.

**Fig 6.**
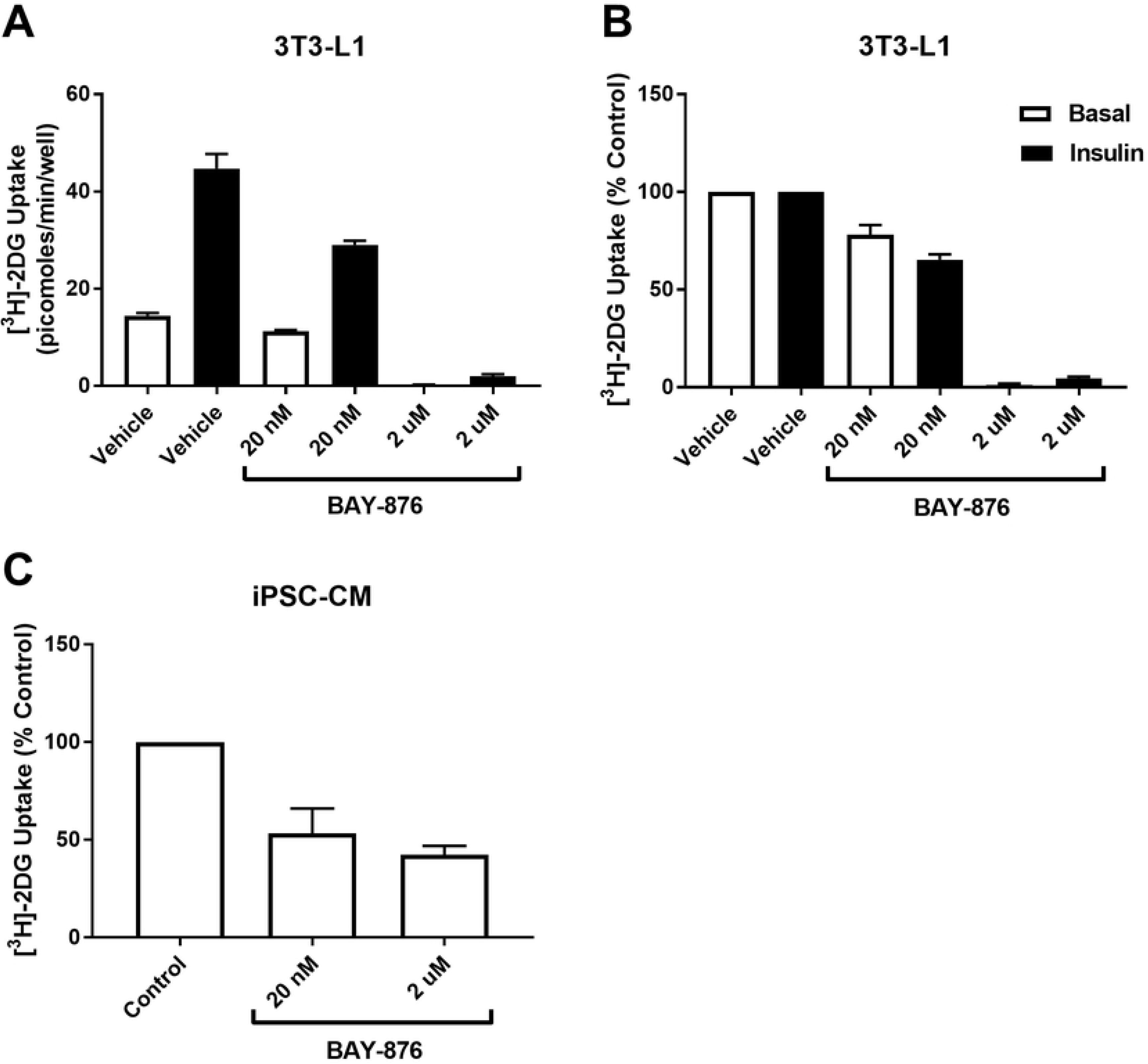
Functional contribution of GLUT1 and GLUT4 to the regulation of cellular glucose uptake in iPSC-CM and 3T3-L1 adipocytes. A: Background corrected basal and insulin stimulated [^3^H]-2-deoxyglucose uptake was assayed in 3T3-L1 adipocytes incubated with 0 [DMSO only], 200 nM or 2 μM BAY-876. Cells were serum starved for 4 hours prior to the assay, incubated with BAY +/− 860 nM insulin for 20 min, and then incubated with the [^3^H]-2-deoxyglucose assay mix for 10 min. Data presented is mean (+S.E.M.) uptake from 3 individual experiments. Statistical analysis was performed with a 2-way ANOVA on raw unadjusted data, and the level of significance was set at P=0.05. B: The same data in panel A presented as the mean (+S.E.M.) percentage difference in values between each experimental group and control basal or control insulin, as appropriate. C: Background corrected basal [^3^H]-2-deoxyglucose uptake was recorded from iPSC-CM that were incubated with 0 [DMSO only], 200 nM or 2 μM BAY-876 and is presented as percentage difference from control. Data are mean (+S.E.M.) uptake for each condition from 3 individual experiments. Statistical analysis was performed with a 2-way ANOVA on raw unadjusted data, and the level of significance was set at P=0.05.

### Attempts to increase iPSC-CM GLUT4 expression

In response to identification of low levels of GLUT4 protein expression in iPSC-CM, several interventions designed to rectify this situation were implemented. During foetal cardiac development there are high levels of GLUT1 and low levels of GLUT4, however this quickly reverses during early postnatal life. The induction of experimental hypothyroidism was found to prevent this switch, thereby implicating the active form of thyroid hormone tri-iodothyronine (T_3_) in the perinatal regulation of cardiac GLUT4 expression [31]. T_3_ has also been used previously in order to induce maturation of other aspects of iPSC-CM physiology, with reported success [33]. Therefore, within the present study iPSC-CM were maintained in medium supplemented with a range of physiologically relevant concentrations of T_3_. However, this failed to significantly alter iPSC-CM GLUT4 expression (supplemental Fig. 2).

Similarly, maturation medium conditioning was implemented, whereby iPSC-CM were maintained in medium containing no or low (1g/L) levels of glucose and were instead forced to rely upon fatty acid metabolism to support ATP regeneration as would be required *in vivo* [23]. However, this resulted in only a modest, non-significant (P>0.05) increase in GLUT4 expression that was overshadowed but a much greater increase in GLUT1 expression (Table 2). This is indicative of a general upregulation of GLUT transporters in response to glucose deprivation, rather than evidence of a specific metabolic maturation effect. Equally, most cell types are highly sensitive to and interact with their environment yet maintaining iPSC-CM for a longer period of time than typically recommended in culture, or in an alternative culture vessel (12 or 24 well plate), had limited impact upon GLUT4 protein expression (data not shown). It was thought that these latter interventions may alter the contractile profile of the cells, which based upon skeletal muscle based studies should act as a stimulus for regulating GLUT4 expression [34].

**Table 2.** Effect of maturation medium conditioning upon iPSC-CM GLUT1 and GLUT4 protein expression.

| Intervention | Components | Ave. Fold change GLUT1 | Ave. Fold change GLUT4 |
| --- | --- | --- | --- |
| **Maturation Medium Condition 1** | 0 mM glucose<br>1x ITS supplement<br>860 nM insulin | 3.6 fold increase | 1.4 fold increase |
| **Maturation Medium Condition 2** | 5.5 mM glucose<br>1 x ITS supplement<br>860 nM insulin | 1.8 fold increase | 0.1 fold decrease |
| **Maturation Medium Condition 3** | 0 mM glucose<br>10% serum<br>172 nM insulin | 6.9 fold increase | 1.8 fold increase |
| **Maturation Medium Condition 4** | 5.5 mM glucose<br>10% serum<br>172 nM insulin | 4.3 fold increase | 1.2 fold increase |
| **Maturation Medium Condition 5** | 0 mM glucose<br>10% serum<br>17.2 nM insulin | 7.9 fold increase | 1.7 fold increase |
| **Maturation Medium Condition 6** | 5.5 mM glucose<br>10% serum<br>17.2 nM insulin | 3.5 fold increase | 1.2 fold increase |
| **Maturation Medium Condition 7** | 0 mM glucose<br>10% serum<br>0 nM insulin | 7.7 fold increase | 1.7 fold increase |
| **Maturation Medium Condition 8** | 5.5 mM glucose<br>10% serum<br>0 nM insulin | 7.2 fold increase | 1.5 fold increase |

Axiogenesis iPSC-CM were maintained in 8 distinct variations of ‘maturation medium’ for 4 days after an initial 3 day recovery period post-plating. All cells were maintained in Dulbecco’s modified eagle medium supplemented with 10 mM HEPES, 2 mM Carnitine, 5 mM Creatine, 5 mM Taurine, 1x non-essential amino acid supplement (ThermoFisher), and 1x linoleic-oleic acid supplement (Sigma). Additional supplements that defined the separate conditions are displayed under ‘components’. GLUT1 and GLUT4 protein expression in iPSC-CM maintained in each of the conditions and control cells was assessed via immunoblotting. Quantification of target protein expression was performed via densitometry and normalised against GAPDH. The data displayed is the mean fold difference in protein expression relative to untreated control cells, generated across 3 independent experiments.

## Discussion

The overarching aim of this study was to assess the potential of iPSC-CM to act as the basis of a novel cellular model of DCM. Primarily, in order to be considered viable candidates, iPSC-CM must exhibit a reproducible and robust insulin-stimulated glucose uptake response. Initial data suggested that this may be possible, based upon the observation that iPSC-CM express and can activate molecules such as Akt and ERK1/2, indicating that key insulin signalling machinery to intermediates involved in stimulating glucose transport are intact. However, we were unable to demonstrate insulin-stimulated glucose transport in iPSC-CMs. A central feature of diabetic physiology is insulin resistance in glucose metabolism. As these cells do not exhibit an insulin response, their utility as a model for DCM must be viewed with caution [23]. This conclusion was reached using cells obtained from 2 separate commercial sources, suggesting that this is not likely to be a limitation unique to iPSC-CM generated from one specific manufacturer.

The wider implications of this dataset for the capacity of iPSC-CM to be used for disease modelling will vary. There are 2 broad approaches to mimicking a disease phenotype with this cell type *in vitro*. First of all, external factors designed to replicate the *in vivo* pathological stimuli may be applied to culture medium and incubated for a sufficient duration in order to induce the desired phenotype. When considering the general importance of metabolism in cellular physiology and phenotype, it could be argued that any model generated via this approach will be of limited clinical relevance. This metabolic limitation is compounded by additional well characterised limitations in iPSC-CM structure and function. In contrast, several studies have reported the spontaneous recapitulation of electrophysiological defects in iPSC-CM generated from patients with known genetic mutations [35,36]. In this instance, when considering a defined pathophysiological process with a clear origin, it could be argued that the poorly representative metabolic phenotype of iPSC-CM (relative to primary cardiomyocytes) may be of lesser importance. However, a key component of drug development is assessment of toxicity, therefore there must be an awareness that iPSC-CM may fail to detect or misrepresent off-target pharmacological effects upon metabolic parameters.

Quantification of GLUT1 and GLUT4 transporter levels in iPSC-CM provided a potential explanation as to why no insulin stimulated glucose uptake response was recorded from these cells. Current understanding is that GLUT4 is the main insulin sensitive glucose transporter [37], therefore very low expression would be expected to limit any insulin response. In contrast GLUT1 is thought to exhibit only a very small degree of insulin sensitivity, achieved via increased general endosomal recycling to the plasma membrane [30]. Consistent with this, Figs.4 and 5 confirms low levels of GLUT4 and high levels of GLUT1 in iPSC-CMs compared to both 3T3-L1 adipocytes and rabbit primary cardiomyocytes. The results of this study also provide new insight into levels of expression of GLUT4 and GLUT1 between these major insulin target tissues. Data obtained using the GLUT-selective inhibitor BAY-876 supports the conclusions drawn regarding the relative functional importance of GLUT1 and GLUT4 in iPSC-CM. In stark contrast to the GLUT4 dependent clear insulin response of 3T3-L1 adipocytes, inhibition of GLUT4 in iPSC-CM had no impact upon 2-deoxyglucose transport. This observation, coupled with the much stronger effect of inhibiting the action of GLUT1 in this cell type, strengthens the suggestion that iPSC-CM may exhibit expression of GLUT transporters (and general physiology) analogous to foetal cardiomyocytes. Additionally, it is of interest that inhibition of the action of GLUT1 and GLUT4 in iPSC-CM only reduced 2-deoxyglucose uptake by approximately 50%, in contrast to 3T3-L1 adipocytes whereby all uptake was essentially abolished. This might suggest that other glucose transporters not classically associated with myocytes may be substantially present in this cell type with notable functional effect.

One intriguing theory that arises from this dataset is that iPSC-CM are already insulin resistant in their baseline condition. This is strengthened by the manufacturer’s recommendation (as for multiple cell lines) that cells are maintained in medium containing high levels of glucose (4.5 g/L or approx. 25 mM). However, the overwhelming reliance upon and strong expression levels of GLUT1 in this cell type more readily suggests that iPSC-CM are indeed just at an earlier stage of cardiac development than adult cells. Therefore, several interventions based upon prior iPSC-CM literature were implemented in an attempt to metabolically mature these cells. However, the results were not indicative of any success. This suggests that whilst simple 2D cell culture approaches are suitable for the differentiation of iPSC-CM, a more advanced approach that better replicates the combination of mechanical, hormonal and metabolic stimuli the heart is exposed to *in vivo* may be required to enhance the maturation of this cell type towards a more clinically relevant cardiac model. Accordingly, the most recent advances in this field have employed novel methods, whereby iPSC-CM were used to form a section of cardiac like tissue and stretched between two termini, allowing simultaneous regulation of loading and contractile mechanics [38]. Alternatively, more direct (but less physiologically relevant) molecular methods could be employed to enhance iPSC-CM GLUT4 protein expression. Alterations in metabolic parameters could realistically drive maturation of other aspects of cellular physiology.

### Conclusion

Overall, we suggest that iPSC-CM are not suitable for use as the basis of a novel cellular model of DCM, due to their lack of insulin stimulated glucose uptake response. This appears to be primarily the result of an immature metabolic phenotype, characterised by a lack of protein expression of the insulin sensitive GLUT4 transporter. Initial attempts to increase iPSC-CM GLUT4 expression were unsuccessful. Development of methods to enhance GLUT4 expression might help realise the significant potential of iPSC-CMs.

## Acknowledgements

This work as supported by a PhD scholarship (FS/14/61/31284) and project grant (PG/18/47/33822) from The British Heart Foundation. Work in GWGs laboratory is supported by grants from Diabetes UK (15/0005246 and 13/0004696). We thank Aileen Rankin for supplying primary cardiomyocytes, Anna Koester and Rachel Livingstone for helpful discussions and Laura Stirrat for expert technical assistance.

## Supplemental Figures

**Supplemental Fig 1. Insulin stimulated [^3^H]-2-deoxyglucose uptake in Axiogenesis iPSC-CM.** Background corrected [^3^H]-2-deoxyglucose uptake was recorded from iPSC-CM, via the protocol detailed in the methods. Cells were insulin stimulated for 30 min prior to incubation with [^3^H]-2-deoxyglucose assay mix for 15 min. Data is displayed as the mean (+S.E.M.) fold change in uptake relative to basal values from 3 representative individual experiments. Statistical testing was performed with a 2-way ANOVA on raw unadjusted data, and the level of significance was set at P=0.05.

**Supplemental Fig 2. GLUT4 protein expression in T_3_ treated iPSC-CM.** iPSC-CM were maintained for 4 days in medium containing 0-13.5 nM T_3_ as indicated. Subsequently, protein lysates were generated and subjected to SDS-PAGE and immunoblotting. Lysates were incubated with antibodies probing for GLUT4 (1:2000, 1% milk, PBS-T) and GAPDH (1:80,000, 1% milk, PBS-T). Quantification was performed via densitometry and the mean (+S.E.M.) expression of GLUT4 (normalised to GAPDH) relative to control for each condition across 3 independent experiments is displayed. Statistical analysis was performed on unadjusted data with a 1-way ANOVA, and the level of significance was set at P=0.05.

## References

1. Kannel WB, Hjortland M, Castelli WP. Role of diabetes in congestive heart failure: the Framingham study. Am J Cardiol. 1974 Jul;34(1):29–34.

2. Boyer JK, Thanigaraj S, Schechtman KB, Pérez JE. Prevalence of ventricular diastolic dysfunction in asymptomatic, normotensive patients with diabetes mellitus. Am J Cardiol. 2004 Apr 1;93(7):870–5.

3. Zahiti BF, Gorani DR, Gashi FB, Gjoka SB, Zahiti LB, Haxhiu BS, et al. Left ventricular diastolic dysfunction in asymptomatic type 2 diabetic patients: detection and evaluation by tissue Doppler imaging. Acta Inform Med. 2013;21(2):120–3.

4. Poulsen MK, Henriksen JE, Dahl J, Johansen A, Gerke O, Vach W, et al. Left Ventricular Diastolic Function in Type 2 Diabetes Mellitus Prevalence and Association With Myocardial and Vascular Disease. 2010;

5. Shang Y, Zhang X, Chen L, Leng W, Lei X, Yang Q, et al. Assessment of Left Ventricular Structural Remodelling in Patients with Diabetic Cardiomyopathy by Cardiovascular Magnetic Resonance. J Diabetes Res. 2016 Jun 22;2016:1–8.

6. Huynh K, McMullen JR, Julius TL, Tan JW, Love JE, Cemerlang N, et al. Cardiac-specific IGF-1 receptor transgenic expression protects against cardiac fibrosis and diastolic dysfunction in a mouse model of diabetic cardiomyopathy. Diabetes. 2010 Jun 9;59(6):1512–20.

7. Schannwell CM, Schneppenheim M, Perings S, Plehn G, Strauer BE. Left Ventricular Diastolic Dysfunction as an Early Manifestation of Diabetic Cardiomyopathy. Cardiology. 2002;98(1–2):33–9.

8. Kobayashi K, Forte TM, Taniguchi S, Ishida BY, Oka K, Chan L. The db/db mouse, a model for diabetic dyslipidemia: molecular characterization and effects of Western diet feeding. Metabolism. 2000 Jan;49(1):22–31.

9. Belke DD, Swanson EA, Dillmann WH. Decreased sarcoplasmic reticulum activity and contractility in diabetic db/db mouse heart. Diabetes. 2004 Dec;53(12):3201–8.

10. Stølen TO, Høydal MA, Kemi OJ, Catalucci D, Ceci M, Aasum E, et al. Interval Training Normalizes Cardiomyocyte Function, Diastolic Ca ^2^+ Control, and SR Ca ^2^+ Release Synchronicity in a Mouse Model of Diabetic Cardiomyopathy. Circ Res. 2009 Sep 11;105(6):527–36.

11. Leto D, Saltiel AR. Regulation of glucose transport by insulin: traffic control of GLUT4. Nat Rev Mol Cell Biol. 2012 Jun 1;13(6):383–96.

12. Peterson LR, Herrero P, Schechtman KB, Racette SB, Waggoner AD, Kisrieva-Ware Z, et al. Effect of Obesity and Insulin Resistance on Myocardial Substrate Metabolism and Efficiency in Young Women. Circulation. 2004 May 11;109(18):2191–6.

13. Scheuermann-Freestone M, Madsen PL, Manners D, Blamire AM, Buckingham RE, Styles P, et al. Abnormal Cardiac and Skeletal Muscle Energy Metabolism in Patients With Type 2 Diabetes. Circulation. 2003 Jun 24;107(24):3040–6.

14. Neubauer S, Horn M, Cramer M, Harre K, Newell JB, Peters W, et al. Myocardial Phosphocreatine-to-ATP Ratio Is a Predictor of Mortality in Patients With Dilated Cardiomyopathy. Circulation. 1997 Oct 7;96(7):2190–6.

15. Belke DD, Larsen TS, Gibbs EM, Severson DL. Altered metabolism causes cardiac dysfunction in perfused hearts from diabetic (*db/db*) mice. Am J Physiol Metab. 2000 Nov;279(5):E1104–13.

16. Semeniuk LM, Kryski AJ, Severson DL. Echocardiographic assessment of cardiac function in diabetic *db/db* and transgenic *db/db*-hGLUT4 mice. Am J Physiol Circ Physiol. 2002 Sep;283(3):H976–82.

17. Boudina S, Bugger H, Sena S, O’Neill BT, Zaha VG, Ilkun O, et al. Contribution of Impaired Myocardial Insulin Signaling to Mitochondrial Dysfunction and Oxidative Stress in the Heart. Circulation. 2009 Mar 10;119(9):1272–83.

18. Takahashi K, Tanabe K, Ohnuki M, Narita M, Ichisaka T, Tomoda K, et al. Induction of Pluripotent Stem Cells from Adult Human Fibroblasts by Defined Factors. Cell. 2007 Nov 30;131(5):861–72.

19. Hwang HS, Kryshtal DO, Feaster TK, Sánchez-Freire V, Zhang J, Kamp TJ, et al. Comparable calcium handling of human iPSC-derived cardiomyocytes generated by multiple laboratories. J Mol Cell Cardiol. 2015 Aug 1;85:79–88.

20. Lee Y-K, Ng K-M, Lai W-H, Chan Y-C, Lau Y-M, Lian Q, et al. Calcium homeostasis in human induced pluripotent stem cell-derived cardiomyocytes. Stem Cell Rev. 2011 Nov;7(4):976–86.

21. Pioner JM, Racca AW, Klaiman JM, Yang K-C, Guan X, Pabon L, et al. Isolation and Mechanical Measurements of Myofibrils from Human Induced Pluripotent Stem Cell-Derived Cardiomyocytes. Stem Cell Reports. 2016 Jun 14;6(6):885–96.

22. Kolanowski TJ, Antos CL, Guan K. Making human cardiomyocytes up to date: Derivation, maturation state and perspectives. Int J Cardiol. 2017 Aug 15;241:379–86.

23. Drawnel FM, Boccardo S, Prummer M, Delobel F, Graff A, Weber M, et al. Disease Modeling and Phenotypic Drug Screening for Diabetic Cardiomyopathy using Human Induced Pluripotent Stem Cells. Cell Rep. 2014 Nov 6;9(3):810–20.

24. Correia C, Koshkin A, Duarte P, Hu D, Teixeira A, Domian I, et al. Distinct carbon sources affect structural and functional maturation of cardiomyocytes derived from human pluripotent stem cells. Sci Rep. 2017 Dec 17;7(1):8590.

25. Sadler JBA, Bryant NJ, Gould GW. Characterization of VAMP isoforms in 3T3-L1 adipocytes: implications for GLUT4 trafficking. Brennwald PJ, editor. Mol Biol Cell. 2015 Feb 1;26(3):530–6.

26. Siebeneicher H, Cleve A, Rehwinkel H, Neuhaus R, Heisler I, Müller T, et al. Identification and Optimization of the First Highly Selective GLUT1 Inhibitor BAY-876. ChemMedChem. 2016 Oct 19;11(20):2261–71.

27. Rajkhowa M, Brett S, Cuthbertson DJ, Lipina C, Ruiz-Alcaraz AJ, Thomas GE, et al. Insulin resistance in polycystic ovary syndrome is associated with defective regulation of ERK1/2 by insulin in skeletal muscle in vivo. Biochem J. 2009;418:665–71.

28. Kramer HF, Witczak CA, Taylor EB, Fujii N, Hirshman MF, Goodyear LJ. AS160 regulates insulin- and contraction-stimulated glucose uptake in mouse skeletal muscle. J Biol Chem. 2006 Oct 20;281(42):31478–85.

29. Graveleau C, Zaha VG, Mohajer A, Banerjee RR, Dudley-Rucker N, Steppan CM, et al. Mouse and human resistins impair glucose transport in primary mouse cardiomyocytes, and oligomerization is required for this biological action. J Biol Chem. 2005 Sep 9;280(36):31679–85.

30. Fischer Y, Thomas J, Sevilla L, Muñoz P, Becker C, Holman G, et al. Insulin-induced recruitment of glucose transporter 4 (GLUT4) and GLUT1 in isolated rat cardiac myocytes. Evidence of the existence of different intracellular GLUT4 vesicle populations. J Biol Chem. 1997 Mar 14;272(11):7085–92.

31. Castelló A, Rodríguez-Manzaneque JC, Camps M, Pérez-Castillo A, Testar X, Palacín M, et al. Perinatal hypothyroidism impairs the normal transition of GLUT4 and GLUT1 glucose transporters from fetal to neonatal levels in heart and brown adipose tissue. Evidence for tissue-specific regulation of GLUT4 expression by thyroid hormone. J Biol Chem. 1994 Feb 25;269(8):5905–12.

32. Calderhead DM, Kitagawa K, Tanner LI, Holman GD, Lienhard GE. Insulin regulation of the two glucose transporters in 3T3-L1 adipocytes. J Biol Chem. 1990 Aug 15;265(23):13801–8.

33. Yang X, Rodriguez M, Pabon L, Fischer KA, Reinecke H, Regnier M, et al. Tri-iodo-l-thyronine promotes the maturation of human cardiomyocytes-derived from induced pluripotent stem cells. J Mol Cell Cardiol. 2014 Jul 1;72:296–304.

34. O’Gorman DJ, Karlsson HKR, McQuaid S, Yousif O, Rahman Y, Gasparro D, et al. Exercise training increases insulin-stimulated glucose disposal and GLUT4 (SLC2A4) protein content in patients with type 2 diabetes. Diabetologia. 2006 Nov 9;49(12):2983–92.

35. Carvajal-Vergara X, Sevilla A, D’Souza SL, Ang Y-S, Schaniel C, Lee D-F, et al. Patient-specific induced pluripotent stem-cell-derived models of LEOPARD syndrome. Nature. 2010 Jun 10;465(7299):808–12.

36. Itzhaki I, Maizels L, Huber I, Zwi-Dantsis L, Caspi O, Winterstern A, et al. Modelling the long QT syndrome with induced pluripotent stem cells. Nature. 2011 Mar 16;471(7337):225–9.

37. Bryant NJ, Gould GW. SNARE Proteins Underpin Insulin-Regulated GLUT4 Traffic. Traffic. 2011 Jun;12(6):657–64.

38. Ronaldson-Bouchard K, Ma SP, Yeager K, Chen T, Song L, Sirabella D, et al. Advanced maturation of human cardiac tissue grown from pluripotent stem cells. Nature. 2018 Apr 4;556(7700):239–43.

